# R-BIND 2.0: An Updated Database of Bioactive RNA-Targeting Small Molecules and Associated RNA Secondary Structures

**DOI:** 10.1101/2022.03.14.484334

**Authors:** Anita Donlic, Emily G. Swanson, Liang-Yuan Chiu, Sarah L. Wicks, Aline Umuhire Juru, Zhengguo Cai, Kamillah Kassam, Chris Laudeman, Bilva G. Sanaba, Andrew Sugarman, Eunseong Han, Blanton S. Tolbert, Amanda E. Hargrove

## Abstract

Discoveries of RNA roles in cellular physiology and pathology are raising the need for new tools that modulate the structure and function of these biomolecules, and small molecules are proving useful. In 2017, we curated the <u>R</u>NA-targeted <u>BI</u>oactive liga<u>N</u>d <u>D</u>atabase (R-BIND) and discovered distinguishing physicochemical properties of RNA-targeting ligands, leading us to propose the existence of an “RNA-privileged” chemical space. Biennial updates of the database and the establishment of a website platform (rbind.chem.duke.edu) have provided new insights and tools to design small molecules based on the analyzed physicochemical and spatial properties. In this report and R-BIND 2.0 update, we refined the curation approach and ligand classification system as well as conducted analyses of RNA structure elements for the first time to identify new targeting strategies. Specifically, we curated and analyzed RNA target structural motifs to determine properties of small molecules that may confer selectivity for distinct RNA secondary and tertiary structures. Additionally, we collected sequences of target structures and incorporated an RNA Structure Search algorithm into the website that outputs small molecules targeting similar motifs without *a priori* secondary structure knowledge. Cheminformatic analyses revealed that, despite the 50% increase in small molecule library size, the distinguishing properties of R-BIND ligands remained significantly different to that of proteins and are therefore still relevant to RNA-targeted probe discovery. Combined, we expect these novel insights and website features to enable rational design of RNA-targeted ligands and to serve as a resource and inspiration for a variety of scientists interested in RNA targeting.

## Introduction

RNA molecules are being recognized as major modulators and therapeutic targets in human diseases ranging from cancer^1^, neurodegenerative^2^ and cardiovascular^3^ diseases as well as viral^4^, bacterial^5^, and fungal^6^ infections. As a consequence, small molecule-based targeting of RNA is gaining increased attention in academia and industry, exemplified by the increasing number of publications with RNA-targeting bioactive ligands (Figure 1A).^7-13^ Recent years serve as both proof and inspiration of the vast potential for successful RNA targeting in the clinic, as shown by the FDA approval of risdiplam for spinal muscular atrophy in August of 2020.^14^ This success represents the first drug targeting a non-ribosomal RNA molecule and thus reminds the community that many unanswered questions remain in the field. Additionally, the growing number of known non-coding RNA functions in various biological systems raises the need for new chemical probes to spatiotemporally investigate the cellular processes regulated by these biomolecules.^15^

**Figure 1:**
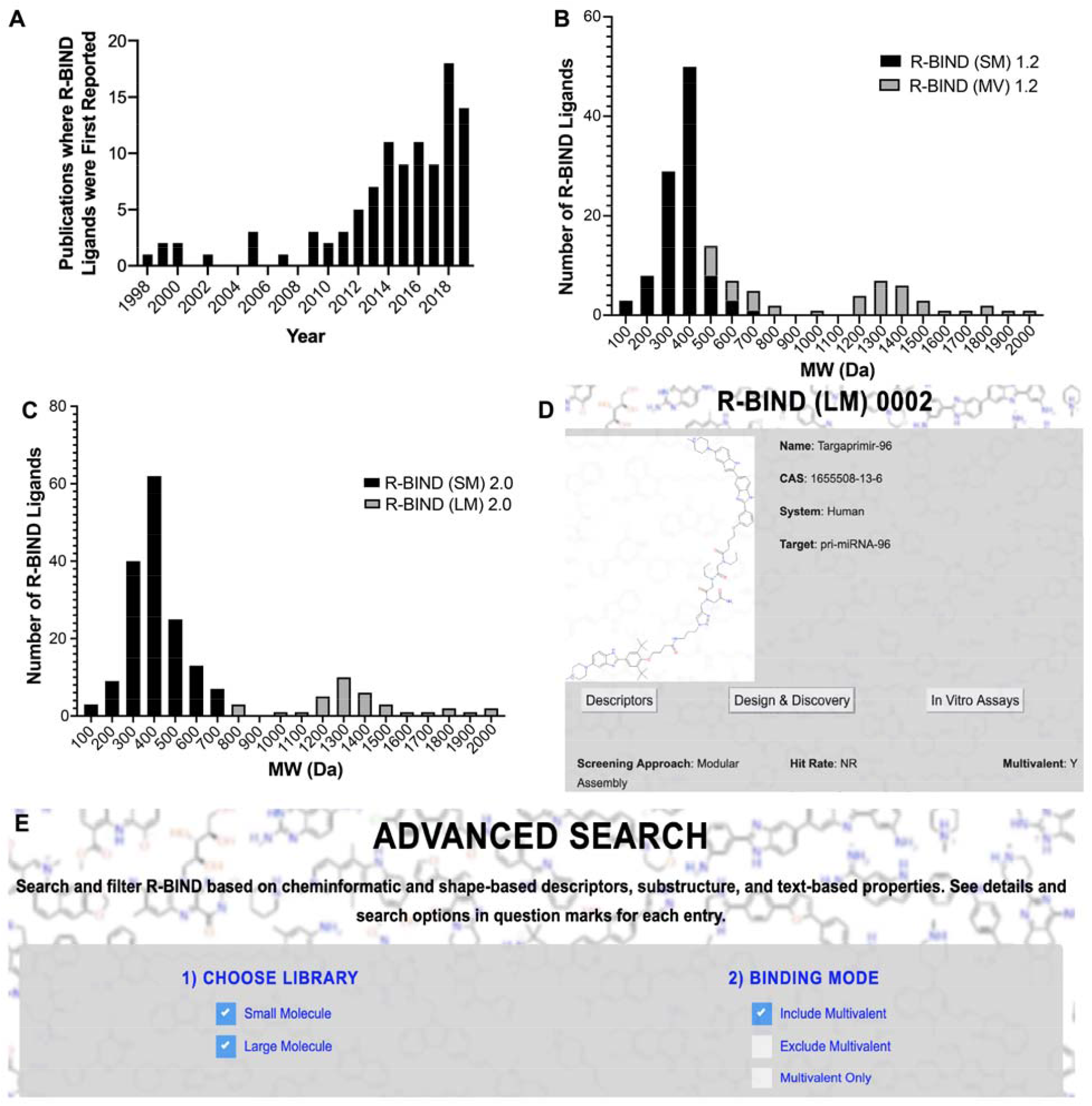
Reclassification of database ligands based on molecular weight (MW) and associated new search features. A) Histogram of articles published reporting ligands that are included in the R-BIND Database. B) Histogram of R-BIND 1.2 classified by SM and MV. C) Histogram of R-BIND 2.0 with new classifications of SM and LM. For histograms, each bin covers 100 Da and the center of the bin is the number listed on the x-axis. For example, bin”200”covers ligands with MW between 150 and 250 Da. D) Website image of new screening approach “Modular Assembly” and “Multivalent: Y” under the Design & Discovery tab in Single Molecule View. E) Website image showing ability to search using multivalency as a criterion under the “Advanced Search.” SM = small molecule; MV = multivalent; LM = large molecule; Y = Yes.

A long-standing hypothesis in the field is that progress towards RNA-targeted probe and drug discovery is hindered by our lack of understanding of the nature of RNA-small molecule interactions, which may be different from those of proteins. Towards elucidating these fundamental principles, our laboratory curated the first database of non-ribosomal RNA-binding ligands to include bioactivity as a criterion for inclusion in 2017, termed the <u>R</u>NA-targeted <u>BI</u>oactive liga<u>N</u>d <u>D</u>atabase (R-BIND).^16^ Its analyses and updates led to the first quantitative comparison and discovery of distinguishing properties for RNA-binding molecules that are different from those of protein-targeting ligands.^16,17^ The existence of this “RNA-privileged chemical space” has been supported by other recent work in the field. For example, similar physicochemical properties were found to be increased in hits in large RNA-targeted screens^18^ and enriching libraries with these properties led to higher propensity for binding to RNA versus protein targets.^19^ Furthermore, recent comprehensive analyses and comparisons of small molecule-bound RNA and protein complexes in the Protein Database (PDB) by Schneekloth and co-workers^20^ and Hargrove and co-workers^21^ revealed differences in pocket properties and interaction types, respectively. The identification of more polar binding pockets in RNA along with the prevalence of stacking and hydrogen bonding interactions in those pockets further support the significance of the enriched molecular recognition parameters identified in R-BIND ligands. Together, these reports corroborate the value of databases such as R-BIND to serve as a reliable resource for designing screening libraries with unique RNA-privileged properties and ultimately expediting the detection of novel probes and drugs.

To make this chemical space and information useful to the community, we recently reported a user-friendly website platform of the R-BIND database (rbind.chem.duke.edu) and offered a number of resources for a diverse community of researchers.^22^ For medicinal chemists, we provided tools to identify novel lead molecules for existing targets via structure-based searches and to design ligands in RNA-privileged space by using a nearest neighbor algorithm. For chemical, molecular, and cell biologists, we extensively curated qualitative information to efficiently select probes and methods for in vitro, cell, and animal experiments. Based on community feedback, and as supported by the established success of structure-based rational design in targeting a variety of RNA structural elements^23^, we envisioned R-BIND users further benefiting from the incorporation of an RNA secondary structure search tool in this update.

In R-BIND 2.0, we conducted an analysis of RNA structure-ligand pairs and identified features that may be privileged for binding particular structural elements. For the field to benefit from these novel insights, we incorporated a feature termed “RNA Structure Search”, an algorithm that allows the user to input an RNA sequence or structure of interest, conducts an RNA structure prediction, and then returns ligands in the database that bind that motif (if any) based on secondary structure and size. We also collected follow-up studies for existing database ligands as well as new reports which were published between June 1^st^, 2018 and December 31^st^, 2019 and found that, despite the small molecule database ligand number increasing by 50% (n = 97 to n = 153), distinguishing cheminformatics and shape-based trends remained the same for R-BIND small molecule database members when compared to protein-targeting ligands. Moreover, we made changes in ligand classification and website information display to more appropriately reflect the set of small molecules as well as highlight distinct strategies of RNA-targeted ligand design. R-BIND 2.0 is complementary to the Disney group’s Inforna database^25^, which compiles data from an in-house two-dimensional combinatorial screen against a panel of RNA secondary structures and requires *a priori* knowledge of the structure or prediction from other prediction platforms. In contrast, R-BIND 2.0 compiles all literature-reported bioactive RNA ligands and their experimental information along with cheminformatic properties and RNA structure-based tools for design and discovery of new ligands.

## Results

### R-BIND 2.0 update includes refined criteria and ligand classification system

R-BIND ligand inclusion criteria are continuously refined and clarified with each update.^16, 17, 22^ In general, we consider chemical probes with molecular weight (MW) <2,000 Da that bind to non-ribosomal RNA targets *in vitro* via non-covalent interactions and have demonstrated activity in cell culture and/or animal models. In this update, we expanded the explicit criteria to state that common nucleic acid intercalators were excluded unless the primary literature report provided experimental evidence that the biological phenotype observed was a result of RNA target binding. The new criteria are outlined in SI Section 1A and in the website “About” and “Contribute” sections.

In addition, we implemented a nomenclature change and re-classification in two ligand libraries. Previously, ligands were separated into a “traditional” small molecule (SM) library (∼ 500 Da) and a multivalent (MV) library. The latter contained larger MW ligands explicitly designed to bind RNA multivalently or those that resemble such ligands and were thus expected to occupy a distinct chemical space. The increased MW of MV library members was expected given their design strategies of combining RNA-binding moieties composed of “traditional” SMs interspaced with specific linkers. All previous cheminformatic analyses and comparisons to FDA-approved drugs were thus conducted with the (SM) library. Throughout previous curations and analyses of the two libraries, however, we noted that while the biggest difference between SM and MV ligands based on physicochemical properties was average molecular weight (MW),^16, 17, 22^ there was overlap between the two libraries. A MW histogram of R-BIND (SM) 1.2 and R-BIND (MV) 1.2 illustrated that the SM ligands MW ranged from 100 to 700 Da, while that of MV ranged from 500 to 2000 Da (Figure 1B).

We reasoned that the exclusion of low-MW MV ligands from the SM group solely based on their design strategy may lead to future biases in the chemical space for RNA-targeted small molecules, which would ideally be defined based on cheminformatics properties and not design strategy. We thus decided to classify two libraries based on a MW cut-off as defined by the largest MW bin in which an R-BIND (SM) 1.2 ligand is located (R-BIND (SM) 0069, MW = 665.91 Da, histogram bin = 700 Da per Figure 1B). Ligands in R-BIND 2.0 were thus classified as follows: small molecules (SM, n = 153) to encompass ligands in bins 700 Da and below, and large molecules (LM, n = 35) for those in bins larger than 700 Da (Figure 1C). To provide information on multivalent ligand design and keep those as searchable features on the website, a new column was added to the “Screening Approach” tab of the “Design and Discovery” section of both libraries to indicate multivalency, and “Modular Assembly” was added as a new Screening Approach category for these ligands (Figure 1D,E). Together, the R-BIND 2.0 update resulted in a total number of 188 R-BIND members, a 25% increase in size compared to 1.2.

### Update 2.0 reveals notable screening trends, novel targets, and advances in discovery strategies

In addition to updating inclusion criteria, we utilize each R-BIND update to note new achievements and emerging strategies in the field. In terms of screening approaches, we observed a significant increase of ligands discovered via focused screens in R-BIND (SM) 2.0 (57%), compared to 36% of the R-BIND (SM) 1.2 library (Figure 2A). In types of primary screens (Figure 2B), we noted the first *in vivo* primary screen in R-BIND 2.0, reported by Chan et. al. in a patent application for DB213 against CAG repeats found in Huntington’s disease.^26^ Specifically, a series of synthesized small molecules were screened in two Drosophila models for polyglutamine-repeat disease. Three compounds were shown to suppress neurodegeneration in a Machado-Joseph disease model and were then tested for CAG-repeat RNA-dependent toxicity in a transgenic Drosophila *DsRed*_*CAG100*_ model. DB213 was shown to rescue the RNA-dependent toxicity in a dose-dependent manner as revealed by a pseudopupil assay in this CAG expansion repeat model but not in control models. Except for this screen, most screens were initially performed *in vitro* rather than through cell-based or computational approaches with bioactivity being assessed in follow-up experiments, consistent with previous updates (Figure 2B). In library types, we noted higher hit rates in focused and synthetic libraries (Figure 2C), which is also consistent with previously described trends.^17^

**Figure 2:**
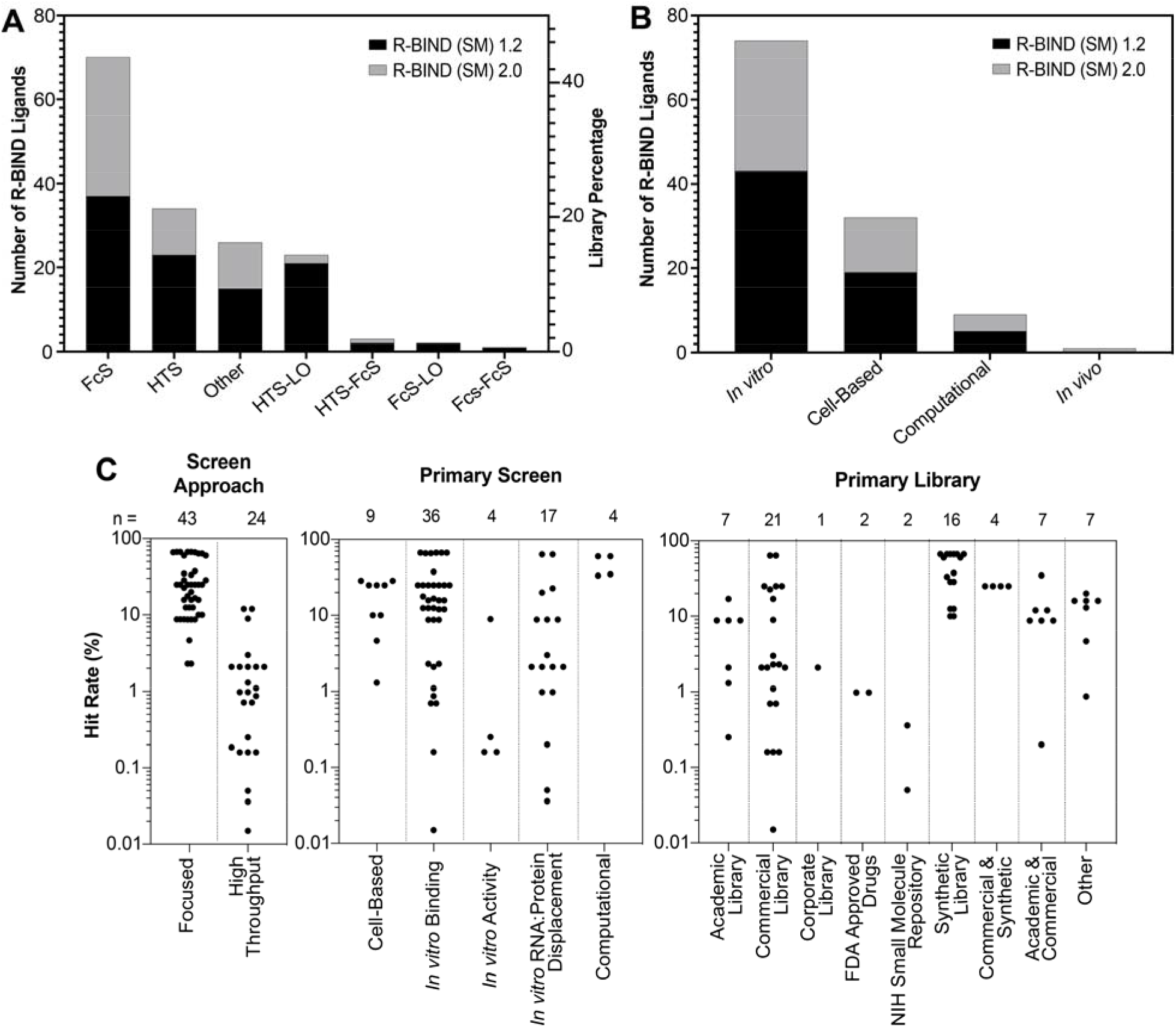
Analysis of screening libraries and approaches utilized in R-BIND (SM) 1.2 and 2.0. A) Number of molecules discovered by various screening approaches by R-BIND (SM) 1.2 and 2.0. B) Number of molecules discovered by primary screen methods in R-BIND (SM) 1.2 and 2.0. C) Hit rates of screens by screening approach, primary screen method, and library source for R-BIND (SM) 2.0. FcS = Focused Screen; HTS = High-Throughput Screen; LO = Lead Optimization.

We also noted increased diversity in targets and discovery strategies. Specifically, advances in long non-coding RNA (lncRNA) targeting were made, as seen by the first reports of bioactive binders for oncogenic MALAT1^27^ (R-BIND (SM) 0113 and 0114) and HOTAIR^28^ (R-BIND (SM) 0137) lncRNAs. Furthermore, an increase in traditionally underrepresented fungal RNA targets was observed with the discovery of first Group II intron binders in Candida Parapsilosis by Pyle and co-workers^6^ (R-BIND (SM) 0121 and 0122), and the first application of differential scanning fluorimetry as a high throughput screen for pre-microRNA 21 inhibitors (R-BIND (SM) 0146, 0147 and 0148) was reported.^29^

Important insights were gained from follow-up studies for previously identified R-BIND ligands, some of which included the addition of new derivatives with the same inferred mode of action. For example, a recent report that outlined structure-activity relationships of antibacterial riboflavin mononucleotide analogs (R-BIND SM 0001 and 0004) led to the addition of R-BIND (SM) 0149 to R-BIND 2.0.^30^ Structural information for small molecule splicing modulators in spinal muscular atrophy determined by NMR (R-BIND (SM) 0095) provided significant insights into the mechanism of action for these ligands,^31^ highlighting the importance of allosteric modulation of RNA-protein complexes in disease.

### Cheminformatic analyses further support existence of RNA-privileged chemical space R-BIND (SM) 1.2 vs. 2.0

Cheminformatic analyses of R-BIND ligands have relied on calculating 20 standard physicochemical properties along with 3D properties using principal moments of inertia (PMI) analysis.^16^ With the addition of new ligands and re-classification of the database for the R-BIND 2.0 update, some of the library averages for the 20 parameters have shifted when compared to the previous (SM) 1.2 update (Table S4). First, an increase in the average molecular weight was observed with the latest update and found to be statistically significantly different per Mann-Whitney *U* analysis in the latest update (*P*-value = 0.02). As a consequence of the MW increase, increases in averages of other parameters were observed as well. For example, accessible surface area (ASA) was found to have a statistically significant increase in R-BIND (SM) 2.0 compared to R-BIND (SM) 1.2 (*P*-value = 0.03). To confirm that these changes were due to the new MW-based library classifications, we conducted a Mann-Whitney *U* Test which showed that if the MW cut-off from R-BIND (SM) 1.2 is applied to the new set of molecules in the R-BIND (SM) 2.0 update, the MW and ASA are no longer significantly different (Table S5). We can thus conclude that these differences were due to reclassification and not a change in the chemical space itself.

To assess the 3D shapes of ligands, we conducted a PMI analysis that classified small molecule shapes as rod-, disc-, or sphere-like. We observed little to no differences between R-BIND (SM) 1.2 and R-BIND (SM) 2.0 updates as assessed by the Kolmogorov-Smirnov test (Table S14), and both libraries remained enriched in rod-like shapes. Taken together, the generally minimal differences in physicochemical and spatial properties, despite another increase in database size, further reinforced the notion that RNA-privileged chemical space exists and can be delineated.

### R-BIND (SM) 2.0 vs. FDA

Before comparing the newly updated R-BIND 2.0 (SM) library to FDA-approved small molecule drugs, our proxy for protein-targeting bioactive ligands, we conducted an update of the FDA library as described in Supporting Information Section 1B. For all cheminformatic analyses, the FDA library used for comparison had a MW cut-off equal to the highest molecular weight ligand in the R-BIND (SM) 2.0 library (R-BIND (SM) 0132 A and B, 705.93 Da), in line with previous comparisons of these two libraries.^16, 17, 22^ In comparing the newly updated libraries, we again conducted a Mann-Whitney *U* analysis and found that 18/20 physicochemical parameters between the FDA and R-BIND 2.0 library had statistically significant differences (Table S6). All structural and molecular recognition parameters that were previously found to be statistically significant between R-BIND (SM) 1.2 and FDA followed the same trend in R-BIND (SM) 2.0 (Figure 3, Table S6 and S7). Some changes in medicinal chemistry properties, as defined by Lipinski’s^32^ and Veber’s^33^ rules, became statistically significant, though the averages do not violate the aforementioned drug-like rules.

**Figure 3:**
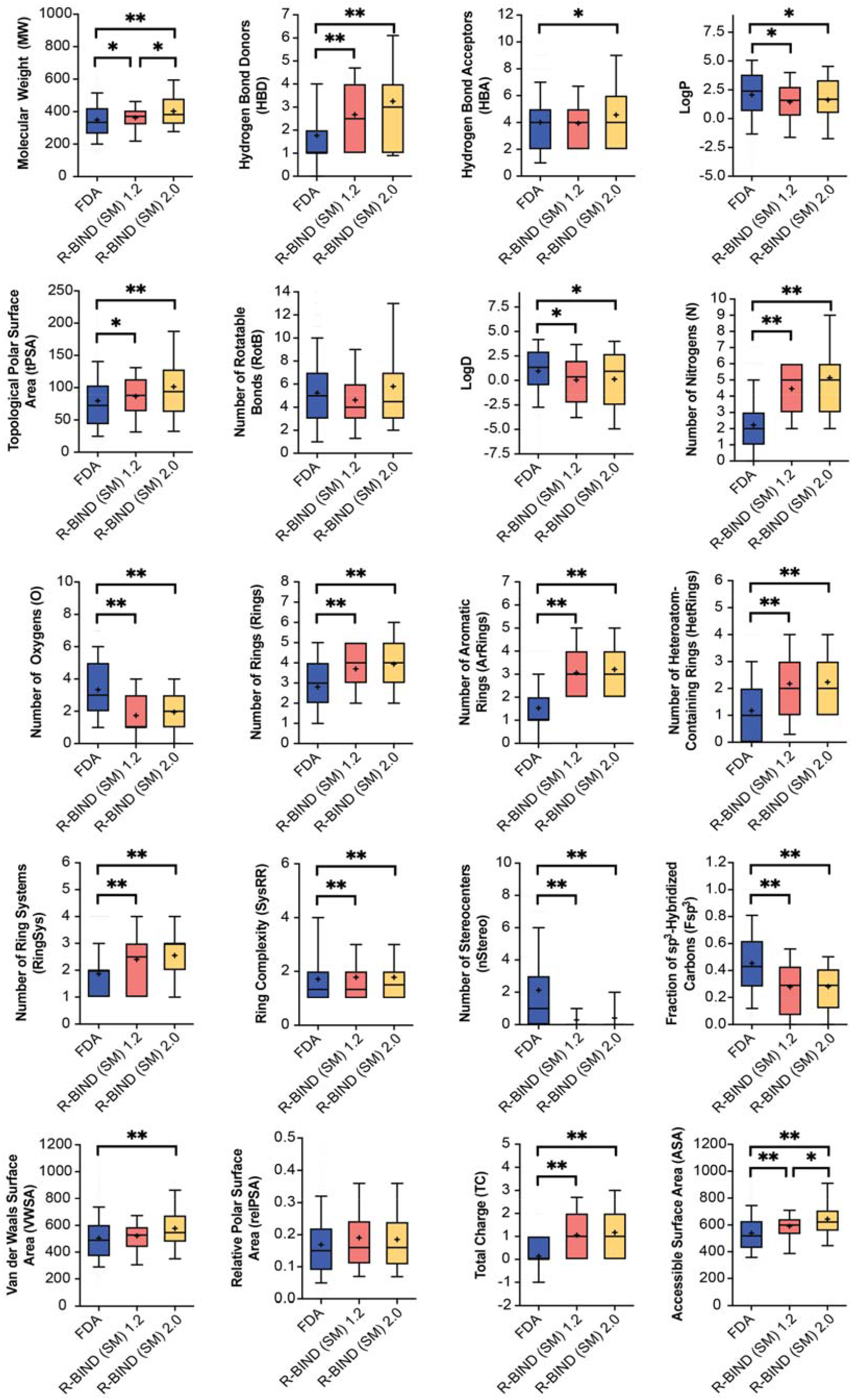
Statistical analysis of cheminformatic parameters between R-BIND (SM) 1.2, 2.0 and the updated FDA library (MW-filtered). The box encompasses 25-75% of variance, while the whiskers describe 10-90%. The mean is indicated by the **+** symbol and the line designates the median value. All comparisons performed using Mann-Whitney U test with statistically significant differences indicated as *P*\* < 0.05, and *P*\*\* < 0.001.

To visualize changes in multi-dimensional chemical space between the FDA, R-BIND (SM) 1.2, and R-BIND (SM) 2.0, we utilized Principal Component Analysis (PCA) as a dimension reduction technique. Notably, the R-BIND library remained in a subset of the space occupied by the FDA library (Figure 4). Several parameters strongly contributed to the first three principal components (Figure S1), which themselves only explain 64% of the total variance (Figure 4D), supporting the need for such multi-dimensional analysis. An expansion of the R-BIND (SM) chemical space was observed as expected from the shifted cheminformatic parameters resulting from the reclassification, and indeed many of the ligands in this expanded space are those that moved from R-BIND (MV) 1.2 to R-BIND (SM) 2.0 (Figure S2). Further, a PCA analysis of R-BIND (SM) and (LM) 2.0 libraries showed unbiased clustering of the two (Figure S3), with major contributors being MW, surface area, and lipophilicity parameters (Table S10, S11 and Figure S4). Lastly, R-BIND 2.0 ligands maintained significantly different enrichment in rod-like character compared to the FDA library (Table S14).

**Figure 4:**
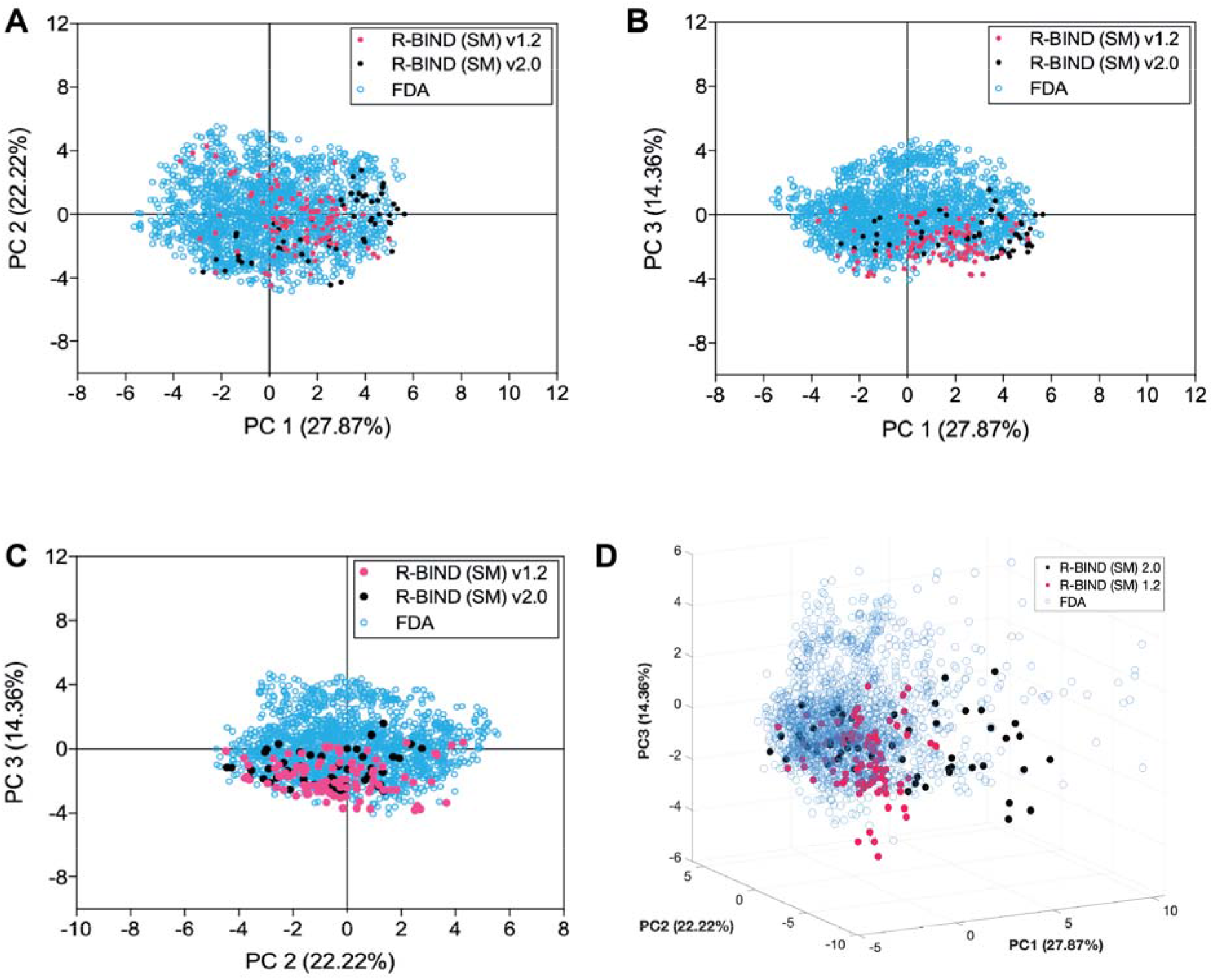
Principal Component Analysis (PCA) plots describing variance between libraries. R-BIND (SM) 2.0 includes only the new ligands between 1.2 and 2.0. Contributions of each parameter are listed in Table S9 and loading plots for the first three principal components are in Figure S1.

### Classification of RNA target structure-ligand pairs reveal differential ligand properties

With the notable increase in the number of ligands in R-BIND (SM) 2.0, we hypothesized that it might be possible to match cheminformatic properties of ligands with their target RNA secondary or tertiary structures when known. This prospect would benefit the existing discovery strategy of rationally designing lead ligands based on RNA sequence and structure similarity. Indeed, Disney and co-workers have demonstrated that their database of RNA motif-small molecule interactions, Inforna^25^, which focuses on affinity for select secondary structures such as loops and bulges can serve to identify binders for targets such as micro-RNAs, viral RNAs, expanded repeat RNAs, and others.^34^ Additionally, Zhang and co-workers have curated a database to explore RNA:ligand interactions by RNA structure, allowing for identification of ligands across three databases that bind to RNA secondary structures while scoring the results based on sequence similarities between the input and reported RNA.^24^

We assessed the potential of our database to reveal novel determinants of structure-based selectivity in a *bioactive* setting for a comprehensive set of literature-reported ligands. Towards this end, linear discriminant analysis (LDA) was performed with the 20 cheminformatics parameters and classification of the SM ligands based on five RNA structural target classes (bulges, G-quadruplexes, double-stranded RNA, internal loops, and stem loops, Table S18). These classes were chosen because they each contained at least 3 ligands as well as ligands for at least three distinct RNA targets within that class, which we deemed as the minimum to obtain meaningful insights into a structural class. We found that the first three components explain ∼88% of the variance within the predefined structure classes and showed significant separation (Figure 5A, Table S19). Small molecule properties such as rotatable bonds, surface area, and the number of ring systems had the greatest contribution to the variance (Figure S8). The separation was further validated through the training and cross-validation of the data set. In the training of the data set, around 84% of small molecules were correctly classified by the RNA structure motif that they target (Table S20). Furthermore, in the cross-validation, around 68% of small molecules were classified correctly (Table S21).

**Figure 5:**
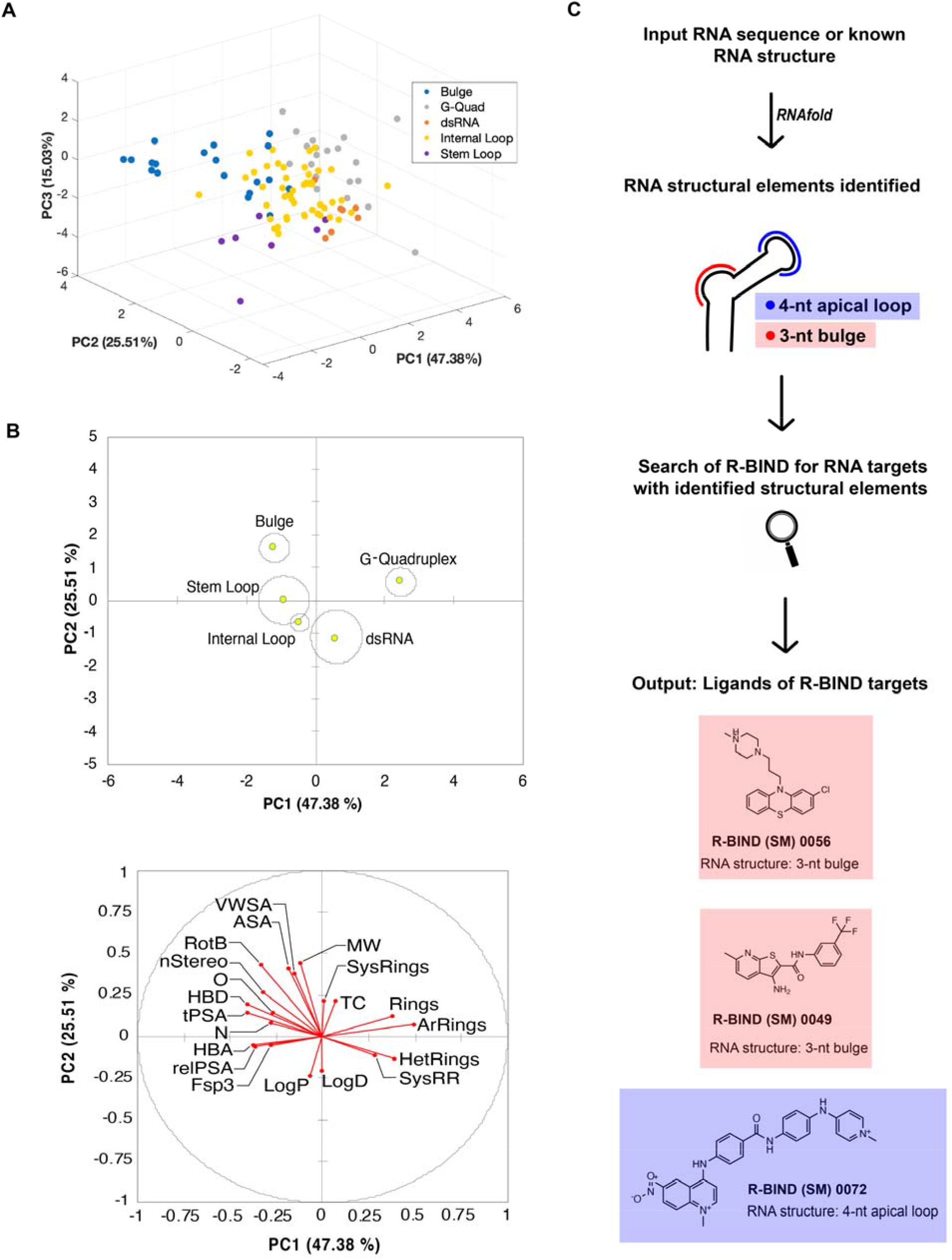
Analysis and implementation of RNA structure targeting using the R-BIND search tools. A) Linear discriminant analysis (LDA) by RNA structure class explained by first 3 principal components (PCs). B) Two dimensional centroid structures and associated loading plot with 20 physicochemical properties. Abbreviations are listed in Figure 3. C) Workflow of newly implemented “RNA Structure Search” on the R-BIND website showing a theoretical input RNA containing a 3-nt bulge and 4-nt apical loop. Briefly, upon inputting a target RNA sequence with or without a dot-bracket notation, the RNAfold algorithm predicts the minimal free energy structure. The structure is inspected for bulge, internal loop, or apical loop structural elements of any size, which is then compared to the structural elements targeted by compounds in the database. The output represents all compounds in R-BIND that bind structures of the same size and motif.

A closer look at two-dimensional centroids and the corresponding loading plot revealed several physicochemical properties that contributed to the structure classification (Figure 5B). For example, G-quadruplex-binding ligands have on average almost 5 rings and over 4 aromatic rings, the highest average number of rings and aromatic rings compared to all other structures examined (Table S22). Ligands that bind dsRNA were enriched in lipophilic character, while those that target more flexible and solvent-exposed bulges had a higher MW, number of rotatable bonds, and surface area parameters. Together, these results show the potential to use cheminformatics and RNA structural information in R-BIND to enrich for ligands targeting a given RNA secondary or tertiary structure.

### Website platform contains updated Nearest Neighbor and novel RNA Structure Search algorithm

#### Nearest Neighbor Updates

The change in the chemical space occupied by R-BIND (SM) 2.0 ligands granted an update of the Nearest Neighbor algorithm, which assesses if an input ligand exists in the RNA-privileged chemical space as defined by the 20 physicochemical parameters. The average shortest Euclidean distance in this 20-dimensional space decreased from 2.0444 for R-BIND (SM) 1.2 to 1.7245 for R-BIND (SM) 2.0, suggesting that the distinct chemical space is becoming more populated.

#### RNA Structure Search Feature

To make the discovery of matching RNA structures with SM ligands of use to the community, we sought to incorporate a feature that enables parsing the database by RNA secondary structure to find small molecules that target the same motif in the R-BIND website. We termed this feature “RNA Structure Search.” Briefly, a user can input an RNA sequence or a defined secondary structure in dot-bracket notation. After conducting a structure prediction embedded in the algorithm in the case that only the sequence is provided, the output will list the simple secondary structure motifs found (bulge, apical loop or internal loop), all small molecules that bind motifs of the same size (e.g., 2-nt bulge), and the sequences of the secondary structure elements that the R-BIND ligands bind (Figure 5C). We note that the embedded feature of predicting the secondary structure based on input sequence takes into account only the lowest-energy structure as analyzed by RNAfold. It is well known that alternative or less-populated structures may exist in the dynamic ensemble of an RNA target of interest, and that environmental factors such as temperature and salt concentrations should be considered when assessing the accuracy of the predicted RNA structure.^35-37^ In the future, we plan on incorporating features in which these factors can be controlled, in addition to predicting the likelihood of forming more complex structures such as G-quadruplexes. In casting a wide net with the RNA Structure Search and identifying all molecules that have been reported to bind a broad set of secondary structure motifs, the promise of RNA-targeting ligands will be a possibility for many therapeutically relevant RNA structures.

## Conclusion

While once considered undruggable, RNA molecules are gaining increased consideration as tractable and sometimes even advantageous targets in disease compared to traditionally targeted proteins.^9^ Timely identification of trends in RNA-small molecule discovery, screening approaches, and RNA-privileged chemotypes along with tool development to apply these findings holds great promise in advancing the field. In this work, we demonstrate that updating the R-BIND database continues to provide insights into the progress and novel strategies employed in the RNA targeting community. We find that the field still embraces focused screens as a screening strategy while introducing innovative high-throughput assays as well as *in vivo* discovery studies. Further, notable progress has been made both in terms of new targets such as long non-coding RNAs, and in the disease phenotypes modulated such as those in Huntington’s disease and fungal infections. Together, insights gained from R-BIND 2.0 confirm that significant progress has been made in the field, and that this trend will continue with years to come.

The analysis of potential changes in physicochemical and spatial properties of ligands in the database continues to define and analyze RNA-privileged chemical space to aid future work. In the R-BIND 2.0 update, addition of new ligands did not change the unique physicochemical properties or rod-like shape preferences found in RNA-targeting small molecules compared to protein-binding ligands. This further supports the hypothesis of the existence of a unique RNA-targeting chemical space. In addition, the increase in library size allowed for insights into physicochemical properties that may distinguish ligands that target specific RNA secondary structures. With advances in pattern recognition^38^- and machine learning^39^-based applications in small molecule modulation of RNA structure, the emergence of initial predictive parameters for structural classes analyzed herein represents an exciting new direction for the field and one that we will continue to explore. One immediate direction is the incorporation of additional cheminformatics parameters such as those used in our recent work for establishing quantitative structure activity relationships.^40^

To increase the utility of the database website and additionally benefit researchers, the website has been updated to incorporate more search features. The new “RNA structure search” includes the ability to input an RNA target of interest and parse the database for similar RNA structural elements. During the preparation of this manuscript, a database termed “RNALigands” from Zhang and co-workers was published,^24^ consisting of R-BIND 1.2, Inforna, and PDB RNA:ligand pairs and offering similar RNA structure-based analyses, highlighting interest in the field to better understand and explore interactions based on RNA target structures. We note, however, that the structure search implemented herein directly queries the R-BIND 2.0 database for bulges, internal loops and apical loops, which were verified as the secondary structures with enough ligands in the database to yield meaningful results. We are confident that these resources can expedite structure-guided rational design of small molecules and aid the exploration of novel RNA targets, including those recently identified in SARS-CoV-2.^24, 41, 42^ Future R-BIND updates and the concurrent increase in RNA structural information will enable synergistic insights that will propel the RNA targeting field to new heights.

## Methods

### Cheminformatics calculations

The non-corrected (NC) SMILES strings for all R-BIND ligands were batch-processed and corrected to their major protonation and tautomer state (pH = 7.4) with ChemAxon calculator plugins (20.8.2). Next, the 20 cheminformatic parameters were calculated using ChemAxon Chemical Terms Evaluator (Marvin 20.8.2, 2020, http://www.chemaxon.com). To compare different libraries, independent two-group Mann Whitney U tests were performed in R software (4.0.0, 2020). The rationale for the selection of cheminformatics parameters as well as descriptions and chemical terms evaluator expressions for all 20 parameters were previously published.^16,22^

### Principal Component Analysis (PCA)

All 20 cheminformatic parameters were normalized to the average and standard deviation of the libraries analyzed as described previously.^16^ The normalized data was used to perform Principal component analysis (PCA) in XLSTAT-Student (version 2019.3.2.61793).

### Principal Moments of Inertia (PMI) Analysis

Calculations were conducted as described previously,^16^ for the R-BIND (SM) v1.2, R-BIND (SM) v2.0, and the FDA 2020 library with the molecular weight restriction (n = 1,834). Protonation- and tautomer-corrected SMILES codes from SI Section 3 were utilized for all libraries. Specifically, the Molecular Operating Environment software was used (MOE, version 2019.01) to set up a conformational search utilizing input parameters listed in Table S12, using a stochastic method with the MMFF94 force field and generalized Born solvation model. The following options were checked: calculate force field partial charges and hydrogens. The normalized (npr1 and npr2) PMI vectors were calculated for each conformation after the search was complete, and the values were Boltzmann-averaged to result in a single coordinate. Details regarding the energy window rationale and Boltzmann average calculation was previously published.^16^ The triangular graph shown in Figure S5 was constructed by plotting the resulting npr1 and npr2 coordinates as vertices of rod- (0,1), sphere- (1,1), and disc-like (0.5,0.5) shapes. The Euclidean distance of each ligand coordinate to each vertex was calculated and ordered from smallest to largest to plot cumulative distance distributions (Figure S6). A two-sided, two-sample Kolmogorov-Smirnov test was conducted to assess the statistical significance of the plotted differences using R statistical software v.3.4.3 (2017) and the results are listed in Table S14. To analyze the shape populations of the libraries, the large triangular plot was partitioned into four (Table S15) or sixteen (Table S16) equal-sized sub-triangles using cell-based partitioning. The triangle partition figures and script details were previously published.^22,16^

### Linear Discriminant Analysis (LDA)

All 20 cheminformatic parameters were used to perform linear discriminant analysis (LDA) in XLSTAT-Student (version 2019.3.2.61793). Only RNA structure classes with 5 or more ligands in R-BIND (SM) 2.0 were included in the LDA analysis (internal loop, bulge, stem loop, g-quadruplex, and dsRNA).

### RNA Structure Search Algorithm

The in-house-written Python algorithm (version 3.7.4) was made to search the unique secondary structure(s) in the user-input connecting (.ct) file, and outputs R-BIND ligands that target the detected secondary structure(s). Details on the algorithm construction rationale and its stepwise workflow can be found in SI Section 7 and Figure S8.

## Supporting information

RBIND_v2.0_A

RBIND_v2.0_B

## Acknowledgments

We are grateful to the members of Hargrove and Tolbert labs for helpful discussions. We thank William Day and Michael Peterson for essential assistance and help with R-BIND website maintenance and addition of new features. TOC graphic was made with BioRender.

## Author contributions

Conceptualization: A.E.H., B.S.T., and A.D.. Database curation and data collection: A.D., E.S., S.L.W., A.U.J., Z.C., K.K., and C.L.. Cheminformatics, principal moments of inertia, principal component, linear discriminant, and nearest neighbor analyses: E.S., S.L.W., and Z.C.. RNA Structure Search algorithm: L-Y C. and A.A.K.. Website integration: B.G.S. and E.H.. Supervision: A.E.H. and B.S.T.. Writing (original draft): A.D., E.S., and A.E.H.. Manuscript editing: All.

