## Supporting Information for "R-BIND 2.0: An Updated Database of Bioactive RNA-Targeting Small Molecules and Associated RNA Secondary Structures"

### 1.Database Curation Criteria

#### A. R-BIND Curation Criteria

Previous updates of the R-BIND database included the following:

- 1.0, which included ligands reported prior to December 31st, 2016[1];
- 1.1, which included ligands reported prior to May 31st, 2017[2];
- 1.2, which included ligands report prior to June 1st, 2018[3];

Manual search to identify potential candidates were conducted as previously described.[3] The current update, 2.0, resulted in the addition of 47 new ligands, including 41 small molecules (SM) and 6 large molecules (LM). A single ligand, R-BIND (SM) 0046, was previously in the database as targeting the HIV-1 Frameshift Site,[4, 5] and was reported to target Huntington's Disease r(CAG) repeats in this update.[6] This ligand was therefore listed as R-BIND (SM) 0046A for the former target report, and R-BIND (SM) 0046B for the latter report. Unique qualitative information specific to each target and associated report is listed in the database. Additionally, a new ligand, R-BIND (SM) 0132, was found to target two distinct pri-miRNA molecules in the same report.[7] Similarly, this ligand was listed as –A and –B for the two different targets, with unique qualitative information available for each target.

To be considered an R-BIND ligand, the molecule(s) followed these guidelines:

- have activity in cell culture and/or animal models;
- not be an aminoglycoside, peptide, oligonucleotide, or known nucleic acid intercalator;
- have evidence of binding to the target *in vitro*;
- have a molecular weight < 2,000 Da;
- target RNA through only non-covalent interactions;
- targeted a non-ribosomal RNA; and
- be highlighted by the author in the manuscript text as being particularly potent or having a notable activity cut-off.

Exceptions for individual criteria were made on a case-by-case basis. For example, R-BIND (LM) 0035 with MW = 2015.63 Da meeting all other criteria was included.[7] R-BIND (SM) 0112 was included despite being a known DNA intercalator, as authors provided experimental evidence for DNA binding not being responsible for the biological phenotype observed.[8] With that same reasoning, R-BIND (SM) 0120 was included, despite being an analog of the DNA intercalator ellipticine; R-BIND (SM) 0118 and 0119 from the same report were included as well because they are analogs of 0120.[9] Finally, two bioactive ligands were included in the update even though there was no evidence of *in vitro* RNA binding, because existing analogs in the database have demonstrated *in vitro* RNA binding. These include R-BIND (SM) 0149,[10] an analog of R-BIND (SM) 0001,[11] and R-BIND (SM) 0150,[12] an analog of R-BIND (SM) 0092.[13]

#### B. FDA Curation Criteria

FDA Approved chemicals were downloaded via DrugBank[14] (v5.1.6, released 04-22-2020) on May 13, 2020. The DrugBank list was filtered to remove molecules that contained the following: <3 carbon atoms (96), metal complexes (112), polymers and oligomers (51), mixtures of unknown composition and ratios (70), inactive ingredients (27), excipients (25), contrast/imaging diagnostic agents or dyes (19), surfactants (16), food coloring (10), preservatives (6), fragrances (5), sweeteners (3), chelating agents (2), and sanitizers (1). At this point, cocktails were split into individual molecules and filtered using the same criteria to remove molecules. Cheminformatics were then performed on this library and the database was filtered by molecular weight (MW) and those molecules with a MW >2,000 Da were removed. For this update, there were a total of 2022 molecules. For statistical comparisons (i.e. Mann-Whitney *U* test), Principal Component Analyses (PCA), and Principal Moments of Inertia (PMI) analyses, the MW cut-off of smallest (140 Da) and largest (706 Da) R-BIND v2.0 (SM) members were applied to the FDA library. This yielded 1,834 molecules for analyses and comparisons to R-BIND (SM).

### 2. R-BIND Data Collection

Information for all ligands in the database is divided into two Excel spreadsheets: "RBIND_v2.0_A.xls", with Overview and Design & Discovery Information as well as cheminformatics information, and "RBIND_v2.0_B.xls", with qualitative experimental information. Both spreadsheets contain separate tabs for SM and LM ligands. To reflect the change in ligand classification system from SM and MV to SM and LM, a column with the previous database ID was added next to the new ID for all ligands in both spreadsheets.

#### A. Spreadsheet RBIND_v2.0_A.xls

The spreadsheet contains five sections as described in the first R-BIND database and website report[3], namely Overview, Target, Quantitative Data, Design & Discovery, and Literature. As a result of the new classification system, a column with the previous database ID a new column "Multivalent" was added to the Design & Discovery section. In the "Multivalent" column, "Y" designates R-BIND ligands that were validated as multivalent binders by the authors, thereby eliminating uncertainty from designating ligands as multivalent unless shown as such in the primary article. The Design & Discovery information is now included for both SM and LM tabs.

The Target Section contains information about the RNA System, Class, Subclass, and Region as listed in SI Table 1. Due to the development of the RNA structure algorithm for the database, the Target section contains two new columns listed next to the RNA Secondary Structure (Binding Location), if known: Secondary Structure Size, where the designation indicates number of nucleotides involved in the structure (e.g. 2,2 indicates a 2-by-2 internal loop); and Secondary Structure Sequence, where the particular nucleotides involved in the structure are listed. In both columns, the number and sequence of nucleotides are listed 5' to 3'. This information is listed in SI Table 2.

**Table S1.** List of R-BIND 2.0 RNA Target systems, classes, subclasses and regions.

| System | RNA Class | RNA Subclass | RNA Region |
|---|---|---|---|
| Bacteria | lncRNA | Expanded Repeats | 5' UTR |
| Fungus | miRNA | Frameshift Site | 5' End |
| Human | mRNA | Heat-Shock Regulatory Element | 3' UTR |
| Virus | mt-mRNA | Internal Ribosome Entry Site (IRES) | 5'/3' UTR |
| | snRNA | pre-miRNA | Exon 1 |
| | vRNA | pre-mRNA | Intron |
| | | Promoter | Nuclease processing site |
| | | Psi Packing Domain | Open reading frames |
| | | Rev Response Element (RRE) | Splice site |
| | | Riboswitch | |
| | | Ribozyme | |
| | | Splicing Complex | |
| | | Trans-Activation Response Element (TAR) | |

**Table S2.** R-BIND 2.0 RNA Target secondary structure names, sizes and sequences. All sizes and sequences are designated as 5'-3'.

| RNA Secondary Structure (Binding Location) | Secondary Structure Size | Secondary Structure Sequence |
|---|---|---|
| Apical Loop | 1 | AACU,GAGC |
| Apical Loop, Bulge | 3 | A |
| Bulge | 4 | A,A |
| Double Stranded RNA (dsRNA) | 5 | AA,U |
| Double Stranded RNA (dsRNA) and ESE2 Motif | 6 | AACUA |
| G-Quadruplex | 1,1 | C,C |
| Internal Loop | 1,1 & 1,1 | C,U |
| Pseudoknot | 2,1 | CUGGGA |
| Stem-Loop | 2,2 | CUGGGA,UCU |
| Three-Way Junction | 3,3 | G,G |
| Triple Helix | 4,4 | G,G & U,U |
| | 6,3 | GGAG |
| | | U |
| | | U,C |
| | | U,U |
| | | UC,CU |
| | | UCU |
| | | UCU,UCU |

#### B. Spreadsheet RBIND_v2.0_B.xls

Spreadsheet B is structured and itemized as described in the original database website report, and details about the individual column descriptions can be found in that report (DOI: 10.1021/acschembio.9b00631.) All current in vitro, in cellulo, and animal model assay categories are listed in SI Table 3.

**Table S3.** List of all in vitro, in cellulo, and animal assay categories in R-BIND 2.0.

| In vitro | In cellulo | Animal Model |
|---|---|---|
| Activity | Antigen Presentation | Antimicrobial Activity |
| Binding | Antimicrobial Activity | Body Temperature |
| In silico | Antiviral Activity | Body Weight |
| Off-Target Effect | Apoptosis | Brain Penetration |
| On-Target Effect | Cell Cycle Progression | Cardiac Function |
| Permeability | Cell Migration | Cecal Flora Disruption |
| Pharmacokinetics | Cell Morphology | Distribution |
| Protein Displacement | Cell Permeability | DNA Abundance |
| RNA Abundance | Cell Uptake | Enzymatic Activity |
| RNA:Protein Stabilization | Cell Viability | Fecal Flora Disruption |
| Structure Determination | Chemoresistance | Lifespan |
| Translational Activity | Colony Formation | Localization |
| | Cytotoxicity | Motor Behavior |
| | Frameshifting Activity | Motor Neuron Pathology |
| | Global mRNA Abundance | Muscle Morphology |
| | Global Protein Abundance | Necrosis |
| | Global RNA Abundance | Neuromuscular Junction Morphology |
| | Inhibitory Potency | Nuclear Foci Dispersion |
| | Ligand Stability | Pharmacokinetics |
| | Localization | Plasma Levels |
| | Metabolic Activity | Protein Abundance |
| | Metabolite Sythesis Inhibition | RNA Abundance |
| | Nuclear Foci Dispersion | RNA Splicing |
| | Off-Target Effect | Rough Eye Phenotype |
| | On-Target Effect | Toxicity |
| | Organoid Branching Morphogenesis | Tumor Burden |
| | Permeability | |
| | Pharmacological Profiling | |
| | Proliferation | |
| | Protein Abundance | |
| | Replication Inhibition | |
| | Reporter | |
| | Ribosome Loading | |
| | RNA Abundance | |
| | RNA Nuclear Export | |
| | RNA Splicing | |
| | Target Engagement | |
| | Transcriptional Activity | |
| | Transcriptional Inhibition | |
| | Translational Activity | |
| | Transport Activity | |

### 3. Cheminformatics Calculations

**Table S4:** Statistical analysis of 20 cheminformatic parameters calculated for R-BIND (SM) v1.2 and the R-BIND (SM) v2.0.

| Category | Parameter | Means | | *P* Value |
|---|---|---|---|---|
| | | R-BIND (SM) v1.2 | R-BIND (SM) v2.0 | |
| Medicinal Chemistry | MW | 365 | 405 | 0.023 |
| | HBA | 3.92 | 4.61 | 0.118 |
| | HBD | 2.74 | 3.21 | 0.738 |
| | LogP | 1.45 | 1.66 | 0.345 |
| | RotB | 4.76 | 5.79 | 0.095 |
| | tPSA | 88.2 | 102 | 0.432 |
| | LogD | -0.049 | 0.211 | 0.402 |
| Structural | N | 4.51 | 5.13 | 0.455 |
| | O | 1.76 | 1.99 | 0.362 |
| | Rings | 3.71 | 3.96 | 0.230 |
| | ArRings | 3.06 | 3.23 | 0.306 |
| | HetRings | 2.16 | 2.26 | 0.561 |
| | SysRings | 2.39 | 2.57 | 0.340 |
| | SysRR | 1.81 | 1.78 | 0.738 |
| Molecular Complexity & Recognition | Fsp3 | 0.280 | 0.277 | 0.949 |
| | nStereo | 0.311 | 0.409 | 0.570 |
| | ASA | 595 | 646 | 0.030 |
| | relPSA | 0.191 | 0.186 | 0.704 |
| | TC | 1.11 | 1.14 | 0.980 |
| | VWSA | 527 | 578 | 0.053 |

**Table S5:** Statistical analysis of 20 cheminformatic parameters calculated for R-BIND (SM) v1.2 and the R-BIND (SM) v2.0 filtered to only include molecules with R-BIND (SM) v1.2 molecular weight (MW) cut-off (665.91 Da, highest MW in R-BIND v1.2). MW = Molecular Weight, HBA = Hydrogen Bond Acceptors, HBD = Hydrogen Bond Donors, LogP = n-Octanol/Water Partition Coefficient, RotB = Number of Rotatable Bonds, tPSA = Topological Polar Surface Area, logD = n-Octanol/Water Partition Coefficient, N = Number of Nitrogen Atoms, O = Number of Oxygen Atoms, Rings = Number of Rings, ArRings = Number of Aromatic Rings, HetRings = Number of Heteroatom-Containing Rings, SysRings = Number of Ring Systems, SysRR = Ring Complexity, $Fsp^3$ = Fraction of $sp^3$-Hybridized Carbons, nStereo = Number of Sterocenters, ASA = Accessible Surface Area, RelPSA = Relative Polar Surface Area, TC = Total Charge, VWSA = Van der Waals Surface Area.

| Category | Parameter | Means | | P Value |
|---|---|---|---|---|
| | | R-BIND (SM) v1.2 | R-BIND (SM) v2.0 (low MW cutoff) | |
| Medicinal Chemistry | MW | 365 | 395 | 0.059 |
| | HBA | 3.92 | 4.53 | 0.197 |
| | HBD | 2.74 | 3.10 | 1.000 |
| | LogP | 1.45 | 1.70 | 0.315 |
| | RotB | 4.76 | 5.47 | 0.182 |
| | tPSA | 88.2 | 98.0 | 0.662 |
| | LogD | -0.049 | 0.278 | 0.352 |
| Structural | N | 4.51 | 4.99 | 0.791 |
| | O | 1.76 | 1.94 | 0.493 |
| | Rings | 3.71 | 3.92 | 0.359 |
| | ArRings | 3.06 | 3.20 | 0.411 |
| | HetRings | 2.16 | 2.27 | 0.548 |
| | SysRings | 2.39 | 2.55 | 0.362 |
| | SysRR | 1.81 | 1.77 | 0.898 |
| Molecular Complexity & Recognition | Fsp3 | 0.280 | 0.276 | 0.897 |
| | nStereo | 0.311 | 0.378 | 0.843 |
| | ASA | 595 | 631 | 0.077 |
| | relPSA | 0.191 | 0.185 | 0.667 |
| | TC | 1.11 | 1.12 | 0.914 |
| | VWSA | 527 | 564 | 0.123 |

**Table S6:** Statistical analysis of 20 cheminformatic parameters calculated for R-BIND (SM) v2.0 and the FDA 2020 molecular weight-filtered library (n = 1,834).

| Category | Parameter | Means | | P Value |
|---|---|---|---|---|
| | | FDA (2020) | R-BIND (SM) v2.0 | |
| Medicinal Chemistry | MW | 349 | 405 | < 0.001 |
| | HBA | 4.01 | 4.61 | 0.010 |
| | HBD | 1.77 | 3.21 | < 0.001 |
| | LogP | 2.06 | 1.66 | 0.018 |
| | RotB | 5.24 | 5.79 | 0.329 |
| | tPSA | 79.7 | 102 | < 0.001 |
| | LogD | 0.961 | 0.211 | 0.004 |
| Structural | N | 2.22 | 5.13 | < 0.001 |
| | O | 3.33 | 1.99 | < 0.001 |
| | Rings | 2.80 | 3.96 | < 0.001 |
| | ArRings | 1.53 | 3.23 | < 0.001 |
| | HetRings | 1.17 | 2.26 | < 0.001 |
| | SysRings | 1.87 | 2.57 | < 0.001 |
| | SysRR | 1.62 | 1.78 | < 0.001 |
| Molecular Complexity & Recognition | Fsp3 | 0.453 | 0.277 | < 0.001 |
| | nStereo | 2.123 | 0.409 | < 0.001 |
| | ASA | 539 | 646 | < 0.001 |
| | relPSA | 0.169 | 0.186 | 0.100 |
| | TC | 0.136 | 1.14 | < 0.001 |
| | VWSA | 505 | 578 | < 0.001 |

**Table S7:** Statistical analysis of 20 cheminformatic parameters calculated for R-BIND (SM) v1.2 and the FDA 2020 molecular weight-filtered library (n = 1,834).

| | | Means | | |
|---|---|---|---|---|
| Category | Parameter | FDA (2020) | R-BIND (SM) v1.2 | *P* Value |
| Medicinal Chemistry | MW | 349 | 365 | 0.026 |
| | HBA | 4.01 | 3.92 | 0.588 |
| | HBD | 1.77 | 2.74 | < 0.001 |
| | LogP | 2.06 | 1.45 | 0.002 |
| | RotB | 5.24 | 4.76 | 0.222 |
| | tPSA | 79.7 | 88.2 | 0.002 |
| | LogD | 0.961 | -0.049 | < 0.001 |
| Structural | N | 2.22 | 4.51 | < 0.001 |
| | O | 3.33 | 1.76 | < 0.001 |
| | Rings | 2.80 | 3.71 | < 0.001 |
| | ArRings | 1.53 | 3.06 | < 0.001 |
| | HetRings | 1.17 | 2.16 | < 0.001 |
| | SysRings | 1.87 | 2.39 | < 0.001 |
| | SysRR | 1.62 | 1.81 | 0.003 |
| Molecular Complexity & Recognition | Fsp3 | 0.453 | 0.28 | < 0.001 |
| | nStereo | 2.123 | 0.311 | < 0.001 |
| | ASA | 539 | 595 | < 0.001 |
| | relPSA | 0.169 | 0.191 | 0.090 |
| | TC | 0.136 | 1.11 | < 0.001 |
| | VWSA | 505 | 527 | 0.06 |

### 4. Principal Component Analysis

**Table S8.** Eigenvalues of each principal component for analysis of R_BIND (SM) v1.2, v2.0 and FDA (MW filtered, n = 1,834).

| | PC 1 | PC 2 | PC 3 | PC 4 | PC 5 | PC 6 | PC 7 | PC 8 | PC 9 | PC 10 |
|---|---|---|---|---|---|---|---|---|---|---|
| **Eigenvalue** | 5.574 | 4.444 | 2.871 | 1.941 | 1.705 | 0.816 | 0.660 | 0.493 | 0.289 | 0.267 |
| **Variability (%)** | 27.871 | 22.222 | 14.357 | 9.704 | 8.527 | 4.081 | 3.298 | 2.465 | 1.447 | 1.336 |
| **Cumulative %** | 27.871 | 50.093 | 64.450 | 74.154 | 82.681 | 86.762 | 90.060 | 92.526 | 93.973 | 95.308 |

| | PC 11 | PC 12 | PC 13 | PC 14 | PC 15 | PC 16 | PC 17 | PC 18 | PC 19 | PC 20 |
|---|---|---|---|---|---|---|---|---|---|---|
| **Eigenvalue** | 0.212 | 0.146 | 0.125 | 0.112 | 0.095 | 0.083 | 0.065 | 0.050 | 0.028 | 0.022 |
| **Variability (%)** | 1.060 | 0.730 | 0.624 | 0.560 | 0.477 | 0.417 | 0.324 | 0.248 | 0.140 | 0.110 |
| **Cumulative %** | 96.368 | 97.098 | 97.722 | 98.282 | 98.760 | 99.177 | 99.501 | 99.750 | 99.890 | 100.000 |

**Table S9.** Percent contributions of each parameter for each principal component for analysis of R-BIND (SM) v1.2, v2.0 and FDA (MW filtered, n = 1,834).

| | PC 1 | PC 2 | PC 3 | PC 4 | PC 5 |
|---|---|---|---|---|---|
| **MW** | 14.713 | 0.742 | 2.004 | 0.126 | 0.288 |
| **HBA** | 8.421 | 7.956 | 0.100 | 0.172 | 3.408 |
| **HBD** | 2.356 | 3.932 | 0.001 | 0.821 | 15.408 |
| **LogP** | 0.049 | 15.182 | 1.208 | 2.474 | 3.783 |
| **RotB** | 5.342 | 0.051 | 2.601 | 21.855 | 4.212 |
| **tPSA** | 6.927 | 11.864 | 0.111 | 0.411 | 0.640 |
| **logD** | 0.030 | 14.192 | 0.830 | 0.493 | 4.813 |
| **N** | 5.112 | 0.285 | 12.938 | 0.185 | 6.522 |
| **O** | 3.841 | 8.193 | 5.243 | 0.726 | 6.912 |
| **Rings** | 8.146 | 3.268 | 0.033 | 15.451 | 1.456 |
| **ArRings** | 4.373 | 4.296 | 14.365 | 0.027 | 0.447 |
| **HetRings** | 6.312 | 0.023 | 5.549 | 9.547 | 0.246 |
| **SysRings** | 7.895 | 2.610 | 4.751 | 0.196 | 0.001 |
| **SysRR** | 0.963 | 0.700 | 1.619 | 30.731 | 2.811 |
| **Fsp3** | 0.043 | 1.003 | 21.738 | 0.527 | 6.154 |
| **nStereo** | 0.947 | 0.930 | 16.865 | 8.468 | 0.053 |
| **ASA** | 12.035 | 2.275 | 1.191 | 5.559 | 0.388 |
| **relPSA** | 0.511 | 17.768 | 2.817 | 0.073 | 1.195 |
| **TC** | 0.003 | 2.679 | 0.000 | 1.688 | 40.576 |
| **VWSA** | 11.982 | 2.052 | 6.036 | 0.469 | 0.687 |

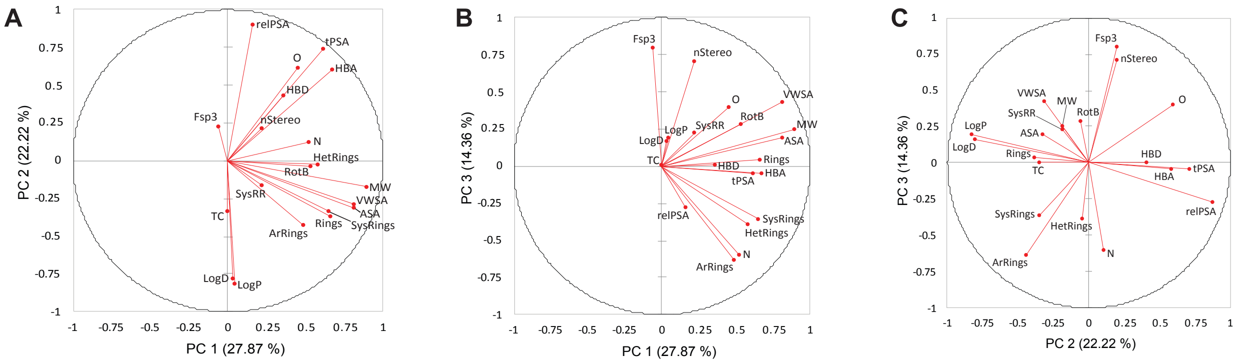

**Figure S1. Loading plots for the first three principal components for analysis of R-BIND v1.2, 2.0 and FDA (MW filtered, n = 1,834).** A) Loading plots for principal component 1 and 2. B) Loading plots for principal component 1 and 3. C) Loading plots for principal component 2 and 3. The magnitude and direction of each vector indicate contribution to the principal components.

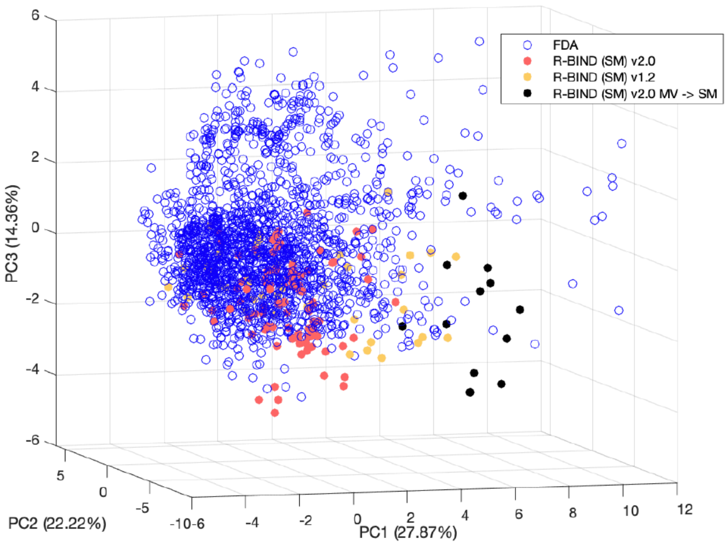

**Figure S2.** 3-Dimensional principal component analysis showing molecules that moved from MV in 1.2 to SM in 2.0 (in black).

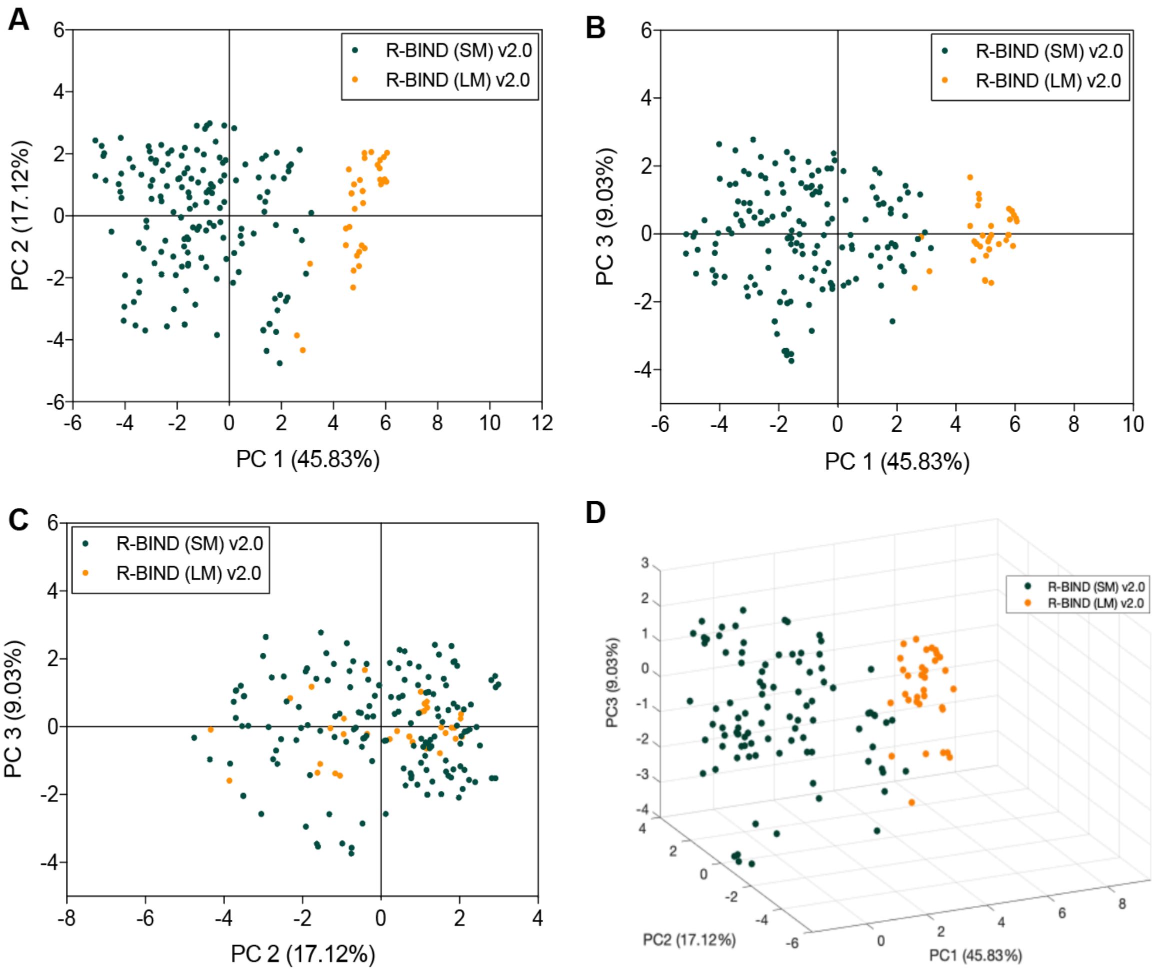

**Figure S3.** Extended principal component analysis (PCA) plots based on the cheminformatics parameters analyzed for R-BIND 2.0 (SM) and (LM) libraries. Each axis represents the principal component (PC) and subsequent percent contribution. A) PCA plot for the first and second principal components. B) PCA plot for the first and third principal components. C) PCA plot for the second and third principal components. D) 3D PCA plot for the first 3 principal components showing the separation of R-BIND 2.0 (SM) ligands from R-BIND 2.0 (LM) ligands.

**Table S10.** Eigenvalues of each principal component for analysis of R_BIND (SM) v2.0 and R-BIND (LM) v2.0.

| | **PC 1** | **PC 2** | **PC 3** | **PC 4** | **PC 5** | **PC 6** | **PC 7** | **PC 8** | **PC 9** | **PC 10** |
|---|---|---|---|---|---|---|---|---|---|---|
| **Eigenvalue** | 9.166 | 3.425 | 1.805 | 1.384 | 1.220 | 0.813 | 0.602 | 0.531 | 0.258 | 0.155 |
| **Variability (%)** | 45.828 | 17.124 | 9.027 | 6.921 | 6.099 | 4.064 | 3.009 | 2.654 | 1.290 | 0.774 |
| **Cumulative %** | 45.828 | 62.952 | 71.979 | 78.899 | 84.998 | 89.062 | 92.071 | 94.726 | 96.015 | 96.789 |

| | **PC 11** | **PC 12** | **PC 13** | **PC 14** | **PC 15** | **PC 16** | **PC 17** | **PC 18** | **PC 19** | **PC 20** |
|---|---|---|---|---|---|---|---|---|---|---|
| **Eigenvalue** | 0.131 | 0.124 | 0.096 | 0.072 | 0.066 | 0.043 | 0.042 | 0.032 | 0.021 | 0.015 |
| **Variability (%)** | 0.653 | 0.621 | 0.480 | 0.358 | 0.330 | 0.217 | 0.210 | 0.161 | 0.106 | 0.075 |
| **Cumulative %** | 97.442 | 98.063 | 98.543 | 98.901 | 99.231 | 99.448 | 99.658 | 99.818 | 99.925 | 100.000 |

**Table S11.** Percent contributions of each parameter for each principal component for analysis of R-BIND (SM) v2.0 and R-BIND (LM) v2.0.

| | **PC 1** | **PC 2** | **PC 3** | **PC 4** | **PC 5** |
|---|---|---|---|---|---|
| **MW** | 9.805 | 0.395 | 0.096 | 0.074 | 0.117 |
| **HBA** | 7.273 | 0.443 | 5.986 | 4.519 | 0.477 |
| **HBD** | 5.347 | 8.269 | 0.147 | 0.940 | 0.125 |
| **LogP** | 0.577 | 15.175 | 0.755 | 1.776 | 9.087 |
| **RotB** | 8.202 | 1.187 | 1.117 | 2.703 | 2.465 |
| **tPSA** | 7.275 | 4.249 | 6.658 | 2.069 | 0.059 |
| **logD** | 0.007 | 21.266 | 2.139 | 0.222 | 8.680 |
| **N** | 8.349 | 1.724 | 0.192 | 0.743 | 0.374 |
| **O** | 4.192 | 0.477 | 1.174 | 21.095 | 5.326 |
| **Rings** | 6.371 | 7.409 | 0.118 | 0.864 | 6.297 |
| **ArRings** | 5.653 | 5.614 | 0.270 | 0.012 | 15.612 |
| **HetRings** | 4.899 | 5.455 | 0.012 | 0.667 | 3.046 |
| **SysRings** | 6.204 | 3.496 | 6.511 | 6.031 | 0.034 |
| **SysRR** | 0.039 | 1.959 | 15.047 | 22.498 | 14.471 |
| **Fsp3** | 1.677 | 0.910 | 17.743 | 0.139 | 23.676 |
| **nStereo** | 2.240 | 0.475 | 6.233 | 17.640 | 7.076 |
| **ASA** | 9.287 | 0.102 | 1.405 | 2.251 | 0.008 |

| | | | | | |
|---|---|---|---|---|---|
| **relPSA** | 0.203 | 13.500 | 20.773 | 5.243 | 0.072 |
| **TC** | 3.036 | 7.836 | 9.775 | 9.591 | 2.929 |
| **VWSA** | 9.365 | 0.059 | 3.848 | 0.921 | 0.068 |

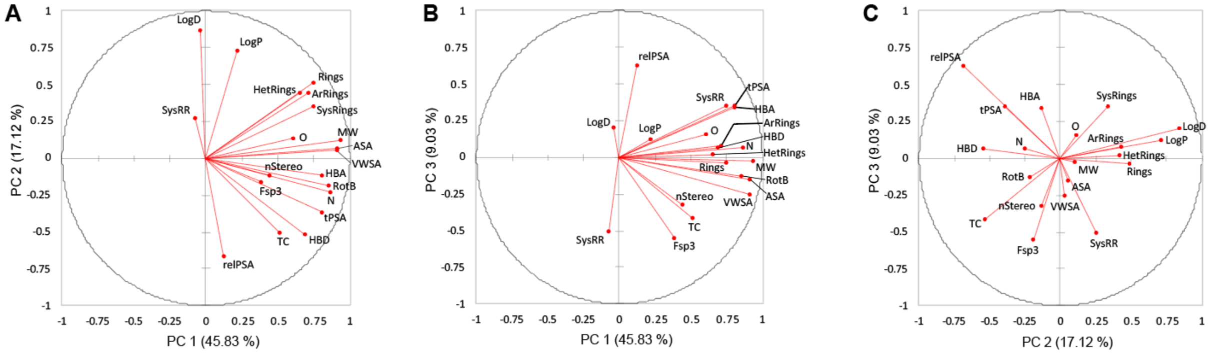

**Figure S4. Loading plots for the first three principal components for analysis of R-BIND (SM) and (LM) v2.0.** A) Loading plots for principal component 1 and 2. B) Loading plots for principal component 1 and 3. C) Loading plots for principal component 2 and 3. The magnitude and direction of each vector indicate contribution to the principal components.

#### 5. Principal Moments of Inertia Calculations

**Table S12**: Input parameters for conformation search

| Parameter | Input |
|---|---|
| Rejection limit | 100 |
| Iteration limit | 10000 |
| RMS gradient | 0.005 |
| MM iteration limit | 500 |
| RMSD limit | 0.15 |
| Energy window | 3 |
| Conformation limit | 10000 |

**Table S13:** Average npr1 and npr2 values for R-BIND (SM) v1.2, v2.0 and MW-filtered FDA 2020 (n = 1,834).

| Library | npr1 | npr2 |
|---|---|---|
| R-BIND (SM) v1.2 | 0.215 | 0.852 |
| R-BIND (SM) v2.0 | 0.239 | 0.849 |
| FDA 2020 (MW) | 0.315 | 0.848 |

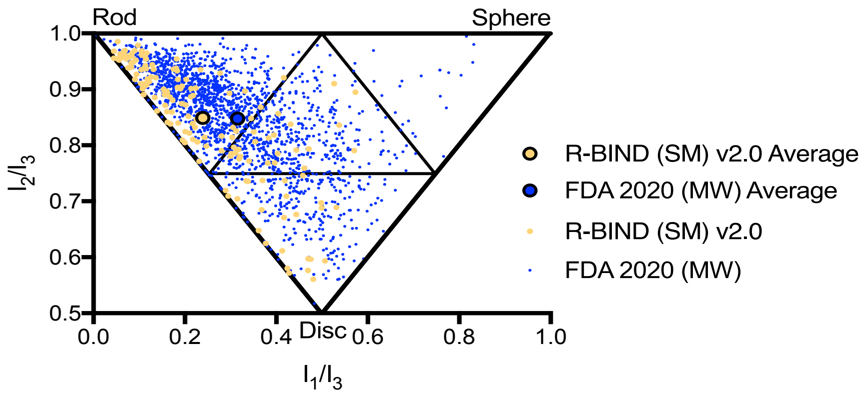

**Figure S5:** Principal Moments of Inertia (PMI) analysis for R-BIND (SM) v2.0 and FDA 2020 library with the MW cut-off. Averages (bolded circles) for both libraries are plotted along with the individual molecules.

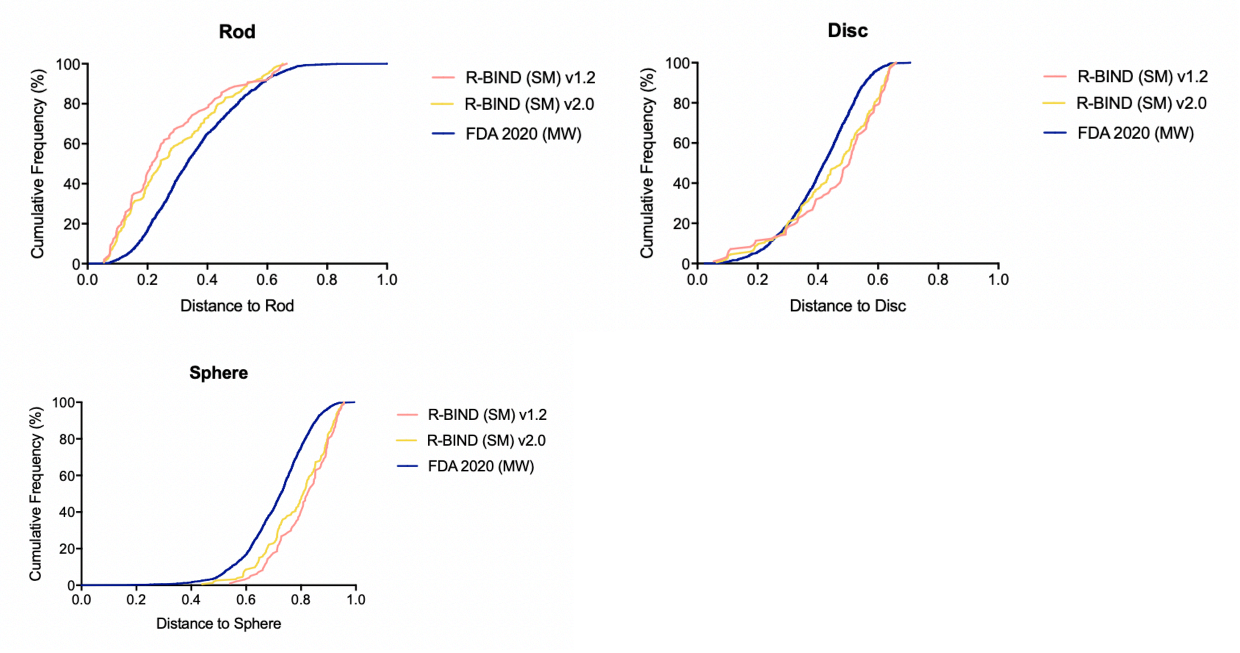

**Figure S6**. Cumulative distributions for distances from rod, disc, and sphere vertices. Graphs were plotted in GraphPad Prism (version 8.4.3 for Mac).

**Table S14.** Statistical comparison from Kolmogorov-Smirnov Test between R-BIND (SM) libraries and FDA library (MW filtered, n = 1,834).

| | ***P-value*** | | |
|---|---|---|---|
| **Library** | **Rod** | **Disc** | **Sphere** |
| R-BIND (SM) v2.0 vs. v1.2 | 0.536 | 0.517 | 0.583 |
| R-BIND (SM) v2.0 vs. FDA 2020 (MW) | <0.001 | <0.001 | <0.001 |

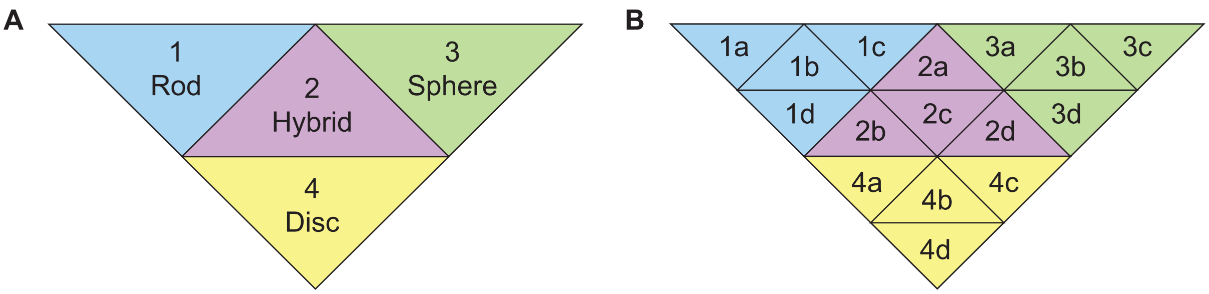

**Figure S7.** Sub-triangle partitioning for A) four- and B) sixteen-triangle partitions used in principal moments of inertia.

**Table S15.** Distribution of small molecules among 4 PMI sub-triangles.

| Library | 1 (Rod) | 2 (Hybrid) | 3 (Sphere) | 4 (Disc) | Total |
|---|---|---|---|---|---|
| R-BIND (SM) v1.2 | 73 | 8 | 0 | 16 | 97 |
| R-BIND (SM) v2.0 | 104 | 24 | 0 | 25 | 153 |
| FDA | 1103 | 441 | 25 | 265 | 1834 |

**Table S16.** Distribution of small molecules among 16 PMI sub-triangles.

| | Rod | | | | Hybrid | | | | Sphere | | | | Disc | | | | |
|---|---|---|---|---|---|---|---|---|---|---|---|---|---|---|---|---|---|
| Library | 1a | 1b | 1c | 1d | 2a | 2b | 2c | 2d | 3a | 3b | 3c | 3d | 4a | 4b | 4c | 4d | Total |
| R-BIND (SM) v1.2 | 36 | 16 | 0 | 21 | 0 | 8 | 0 | 0 | 0 | 0 | 0 | 0 | 5 | 3 | 0 | 8 | 97 |
| R-BIND (SM) v2.0 | 51 | 21 | 1 | 30 | 2 | 17 | 4 | 1 | 0 | 0 | 0 | 0 | 12 | 4 | 0 | 9 | 153 |
| FDA | 260 | 472 | 51 | 320 | 47 | 240 | 112 | 42 | 6 | 8 | 4 | 7 | 115 | 71 | 41 | 38 | 1834 |

**Table S17:** Comparison of small molecules whose triangle assignment based on PMI analysis changed between v1.2 and v2.0 calculations.

| | R-BIND (SM) v1.2 Analysis | | | R-BIND (SM) v2.0 Analysis | | |
|---|---|---|---|---|---|---|
| Molecule ID | npr1 | npr2 | Triangle | npr1 | npr2 | Triangle |
| R-BIND (SM) 0027 | 0.232 | 0.894 | 1b | 0.230 | 0.855 | 1d |
| R-BIND (SM) 0039 | 0.419 | 0.668 | 4b | 0.417 | 0.671 | 4a |
| R-BIND (SM) 0042 | 0.367 | 0.721 | 4a | 0.526 | 0.910 | 2a |
| R-BIND (SM) 0067 | 0.411 | 0.837 | 2b | 0.404 | 0.857 | 2c |
| R-BIND (SM) 0086 | 0.215 | 0.915 | 1b | 0.207 | 0.852 | 1d |
| R-BIND (SM) 0096 | 0.410 | 0.821 | 2b | 0.573 | 0.895 | 2a |

### 6. Linear Discriminant Analysis

**Table S18.** Number of molecules from R-BIND (SM) known to target each structure class represented in the library.

| Structural Class | Number of Small Molecules |
|---|---|
| Internal Loop | 51 |
| Bulge | 20 |
| G-Quadruplex | 20 |
| dsRNA | 7 |
| Stem Loop | 7 |
| Apical Loop | 4 |
| Triple Helix | 2 |
| Pseudoknot | 1 |
| Three-way Junction | 1 |

**Table S19.** Eigenvalues of each principal component for analysis of R-BIND 2.0 RNA structural elements.

| | PC1 | PC2 | PC3 | PC4 |
|---|---|---|---|---|
| Eigenvalue | 1.687 | 0.908 | 0.535 | 0.430 |
| Discrimination (%) | 47.380 | 25.512 | 15.032 | 12.076 |
| Cumulative % | 47.380 | 72.892 | 87.924 | 100.000 |

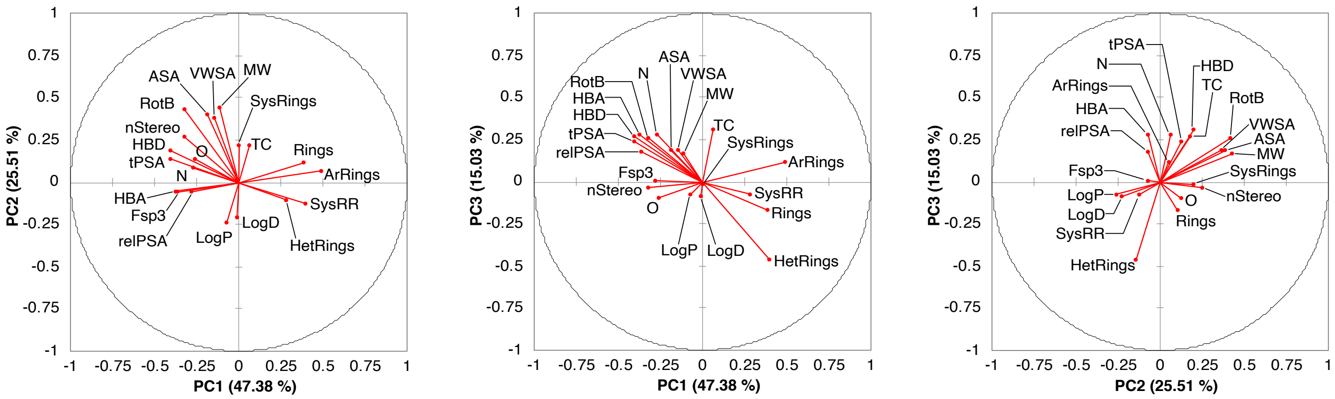

**Figure S8. Loading plots for the first three principal components for analysis of RNA structure targeting.** A) Loading plots for principal component 1 and 2. B) Loading plots for principal component 1 and 3. C) Loading plots for principal component 2 and 3. The magnitude and direction of each vector indicate contribution to the principal components.

**Table S20.** Confusion matrix for the training sample.

| from \ to | Bulge | G-Quadruplex | Internal Loop | Stem Loop | dsRNA | Total | % correct |
|---|---|---|---|---|---|---|---|
| Bulge | 12 | 0 | 8 | 0 | 0 | 20 | 60.00% |
| G-Quadruplex | 0 | 17 | 3 | 0 | 0 | 20 | 85.00% |
| Internal Loop | 1 | 1 | 48 | 1 | 0 | 51 | 94.12% |
| Stem Loop | 0 | 1 | 2 | 4 | 0 | 7 | 57.14% |
| dsRNA | 0 | 0 | 0 | 0 | 7 | 7 | 100.00% |
| Total | 13 | 19 | 61 | 5 | 7 | 105 | 83.81% |

**Table S21.** Confusion matrix for the cross validation.

| from \ to | Bulge | G-Quadruplex | Internal Loop | Stem Loop | dsRNA | Total | % correct |
|---|---|---|---|---|---|---|---|
| Bulge | 11 | 1 | 8 | 0 | 0 | 20 | 55.00% |
| G-Quadruplex | 1 | 13 | 5 | 0 | 1 | 20 | 65.00% |
| Internal Loop | 3 | 5 | 40 | 2 | 1 | 51 | 78.43% |
| Stem Loop | 0 | 1 | 2 | 4 | 0 | 7 | 57.14% |
| dsRNA | 0 | 2 | 2 | 0 | 3 | 7 | 42.86% |
| Total | 15 | 22 | 57 | 6 | 5 | 105 | 67.62% |

**Table S22.** Number of rings and aromatic rings in R-BIND (SM) library showing the enrichment of these groups in g-quadruplex binding small molecules.

| RNA Motif | Average Number of Rings | Average Number of Aromatic Rings |
|---|---|---|
| G-Quadruplex | 4.9 | 4.3 |
| dsRNA | 4.71 | 3.71 |
| Stem Loop | 4.14 | 2.71 |
| Bulge | 4.06 | 3.11 |
| Internal Loop | 3.80 | 3.20 |
| Apical Loop | 3.25 | 3.25 |

### 7. RNA Structure Search

In this RNA secondary structure search algorithm, we aim to provide potential small molecules which bind to an RNA of interest based input the RNA sequence and predicted structure. We adopted an empirical approach by collecting literature reported details of binding sites for R-BIND ligands. In R-BIND 2.0, the highest reported secondary structural features were Internal loops, bulges and G-quadruplexes (Table S18). An algorithm was designed to search secondary structure features (internal loops, bulges and apical loops) based on their shape. We focused on extracting the RNA binding pocket based on the size of loop, such as N-nt bulge, N-nt apical loop and N X M internal loop, where N indicates the size of internal loop in the 5'-end and M indicates the size of internal loop in the 3'-end. For instance, an input RNA sequence that has a 4 X 4 internal loop will output a benzimidazole hit (R-BIND (SM) 0040) from the database, which was reported to bind the 4 X 4 internal loop of the HCV-IRES.[15]

Generally, this algorithm allows users to input their RNA sequence of interest with or without secondary structure information in dot-bracket format. An RNA sequence without secondary structure information will automatically have its secondary structure predicted by RNAfold module,[16] and only the most thermodynamically stable (lowest energy) secondary structure is used for the database search. Both inputs with or without dot-bracket notation will generate a connecting file (.ct file) with the dot-bracket notation of the RNA searched. The stepwise algorithm works as follows (Fig S9):

**Flowchart of RNA structure search algorithm:**

1. Find a target RNA sequence. Ensure that any thymidines were replaced with uridines.
2. Enter target RNA sequence.

3A. If RNA structure is unknown, the system automatically folds target RNA sequence by RNAfold algorithm.

4. The algorithm will output the most stable structure information with connecting file (.ct) format, if there is no experimental data for the secondary structure information of target RNA sequence (such as SHAPE, DMS, NMR or X-ray structural data).

3B. If RNA structure is known, enter target RNA secondary structure information in dot-bracket format.

5. The RNA secondary structure in dot-bracket format from step 4 or 5 is converted into connecting file (.ct) format.
6. The connecting file is analyzed using an in-house-written Python program to extract the internal loops, bulges and/or apical loops. Note, in this algorithm, an N X M internal loop is treated the same as an M X N internal loop.
7. Search R-BIND for matching secondary structures.
8. Output the small molecule(s) which bind to the RNA with the identified secondary structure(s) in the database. While the search is not sequence specific, the sequence composition of the identified structure(s) is also provided to the user as a starting point to optimize RNA binding affinity if it matches an entry in the database.

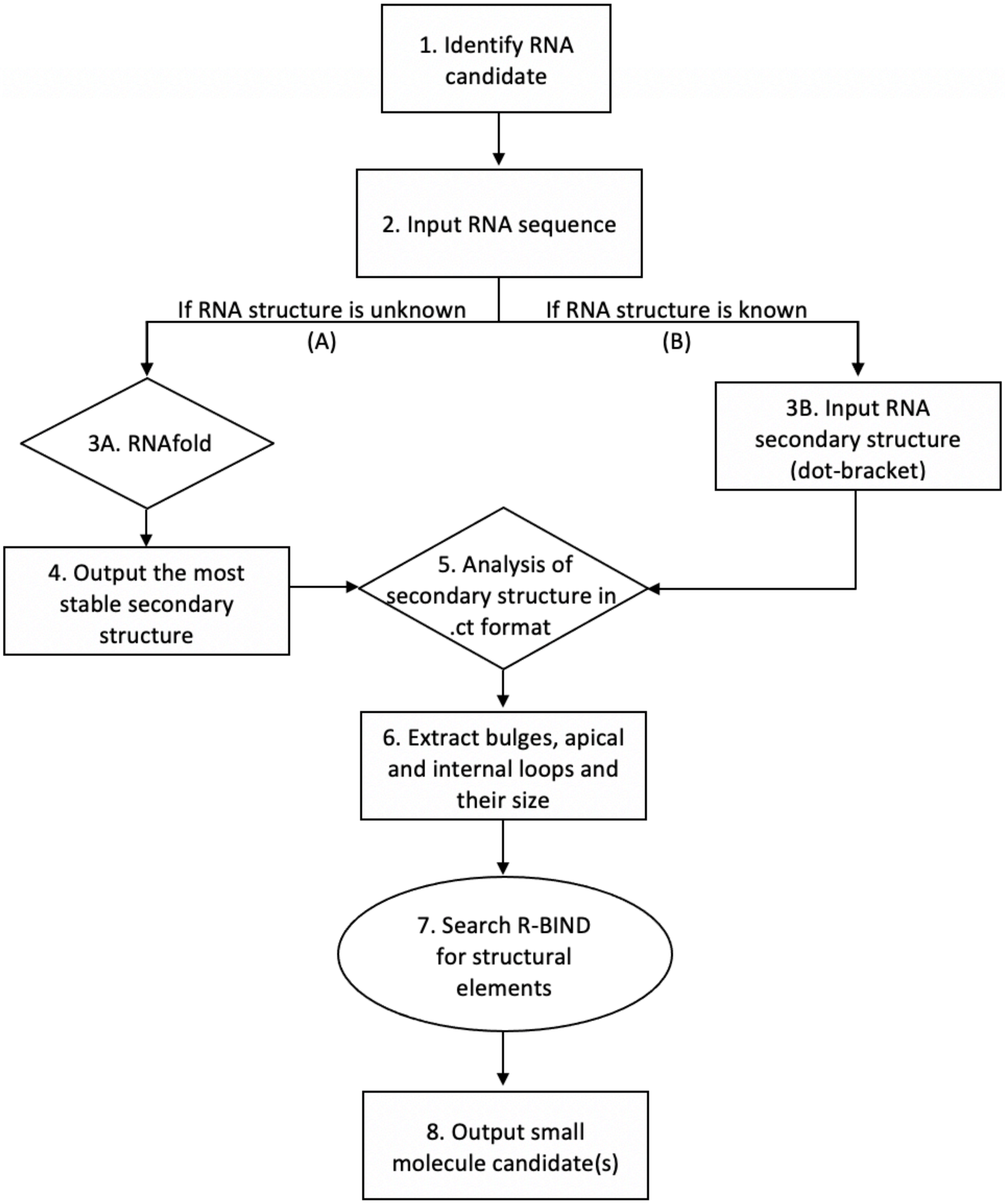

**Figure S8. Flowchart of RNA structure search algorithm.** Route A shows the RNA structure search without a *priori* knowledge of RNA secondary structure. Alternatively, users can choose to input secondary structure information in a dot and bracket format with a *priori* knowledge through route B.
